# Modification of plant regeneration medium decreases the time for recovery of Solanum lycopersicum cultivar M82 stable transgenic lines

**DOI:** 10.1101/046839

**Authors:** Sarika Gupta, Joyce Van Eck

## Abstract

Tomato (Solanum lycopersicum) has rapidly become a valuable model species for a variety of studies including functional genomics. A high-throughput method to obtain transgenic lines sooner than standard methods would greatly advance gene function studies. The goal of this study was to optimize our current transformation method by investigating medium components that would result in a decreased time for recovery of transgenics. For this study, 6-day-old cotyledon explants from Solanum lycopersicum cultivar M82 in vitro-grown seedlings were infected with the *Agrobacterium tumefaciens* strain LBA4404 containing the binary vector pBI121. This vector contains the β-glucuronidase reporter gene and the neomycin phosphotransferase II selectable marker gene that confers resistance to kanamycin. Modification of our standard plant regeneration medium with indole-3-acetic acid (IAA) at concentrations of either 0.05 mg/l or 0.1 mg/l decreased the recovery time for transgenic lines by 6 weeks as compared to our standard medium that contains zeatin as the only plant growth regulator. We observed 50% and 54% transformation efficiency on plant regeneration medium containing 0.05 mg/l and 0.1 mg/l IAA, respectively. Moreover, addition of 1 mg/l IAA to the root induction medium resulted in earlier root development than medium that did not contain IAA. Addition of IAA to the plant regeneration and rooting media did not have any negative effects on plant development. Recovery of transgenic lines in a shorter time results in higher throughput for the introduction of gene constructs and has the potential to decrease the time and resources needed to complete investigations of gene function.

## Introduction

Tomato, *Solanum lycopersicum*, is a member of the *Solanaceae* family, which contains approximately 3,000 plant species and includes some of the most economically important food crops. It is native to South America and was brought to Europe in the 1500s and then to North America in the 1800s (Jones 1998). Tomato is a perennial plant that has two different growth habits, determinate and indeterminate. There are two different market types of tomatoes, fresh market and processing. According to the Agricultural Marketing Resource Center, in 2014 the US dollar value for fresh market tomatoes was 1.14 billion and 1.325 billion for processing types, which are used to make products such as juice, sauces, and ketchup [2]. in addition to being an economically important food crop, tomato is an excellent source of health beneficial nutrients including beta-carotene and lycopene.

Over the years, utilization of tomato as a model plant species has increased because of readily available resources such as mutant populations (Emmanuel and Levy 2002), bioinformatics tools (Bombarely et al. 2011), and a high quality reference genome (Consortium 2012). In addition, since the very first report of *Agrobacterium*-mediated transformation of tomato by McCormick et al. (Mccormick et al. 1986), there have been other reports of successful transformations of different genotypes (Chyi and Phillips 1987; Fillatti et al. 1987; Frary and Earle 1996; Park et al. 2003; Sun et al. 2006; Van Eck et al. 2006) and methods to improve transformation efficiency (Dan et al. 2016). A key aspect for the adoption of a model plant species is the availability of efficient transformation methodology. This was certainly the case for *Arabidopsis*, which is by far the most widely used model for plant research programs (Somerville and Koornneef 2002).

While there are several methods available for plant transformation, A*grobacterium tumefaciens*-mediated transformation has become the most extensively used method (Gelvin 2003; Pitzschke and Hirt 2010). Despite its effectiveness for gene transfer in tomato, there is still need for improvement. Improving methodology to decrease the time from introduction of a gene construct of interest to recovery of stable transgenics would improve the throughput and shorten the timeframe for studies that utilize tomato transgenic lines.

We were interested in finding an approach to decrease the time to obtain transgenic lines of the processing type tomato M82 because this genotype is used for gene function studies in our lab as well as others (Brooks et al. 2014; Xu et al. 2015). We chose to start by investigating supplementation of our standard plant regeneration and rooting media with a growth regulator that had the potential to speed up plant development (Van Eck et al. 2006).

Cytokinins and auxins are important hormones that influence growth and developmental processes in plants. Interactions between cytokinins and auxins have been shown to be necessary for the shoot apex growth (Gupta and Rashotte 2012; Shimizu-Sato et al. 2009). Auxin has also been shown to play a role in the specification of the root apical meristem (Friml et al. 2003; Gupta and Rashotte 2012; Sabatini et al. 1999). The hormonal interactions can be utilized in the area of tissue culture to leverage the presence of the hormones in the medium. In this study, we report the effects of the addition of the auxin, indole-3-acetic acid (IAA) on the recovery time of M82 transgenic lines.

## Materials and methods

### Plant material

Seeds of *Solanum lycopersicum* cv M82 were surface sterilized in 20% (v/v) bleach solution containing Tween-20 for 20 min followed by 3 rinses in sterile water. Seeds were germinated in Magenta GA7 boxes (Caisson Labs, Logan, UT) that contained 50 ml of Murashige and Skoog (MS) (Murashige and Skoog 1962) (Caisson Labs) based medium containing 2.15 g/l MS salts, 100 mg/l myo-inositol, 2 mg/l thiamine, 0.5 mg/l pyridoxine, 0.5 mg/l nicotinic acid, 10 g/l sucrose and 8 g/l Sigma agar (Sigma-Aldrich, St. Louis, MO). Cultures were maintained at 24 °C under a 16h light/8h dark photoperiod at 57 – 65 *u*E m^-2^ s^-1^.

One day prior to infection with *Agrobacterium*, cotyledon explants and feeder layer plates were prepared. Feeder layers were prepared before cutting the explants by dispensing 2 ml of a 1-week-old NT1 suspension culture onto KCMS medium (4.3 g/l MS salts, 100 mg/l myo-inositol, 1.3 mg/l thiamine, 0.2 mg/l 2,4-dichlorophenoxy acetic acid, 200 mg/l KH_2_PO_4_, 0.1 mg/l kinetin, 30 g/l sucrose, 5.2 g/l Agargel (Sigma Aldrich), pH 6.0. The suspension was covered with a sterile 7 cm Whatman filter paper. Explants were excised from 6-day-old seedlings before the first true leaves emerged. To prepare the explants, seedlings were placed on a sterile paper towel moistened with sterile water. Cotyledons were excised at the petioles, cut into approximately 1 cm sections, placed adaxial side down on the KCMS feeder layer plates, and maintained at 24 °C under a 16 h light/8 h dark photoperiod at 57 – 65 *u*E m^-2^ s^-1^.

### Bacterial strain and binary vector

Electroporation was used to introduce the pBI121 vector (Chen et al., 2003) into the *Agrobacterium tumefaciens* strain LBA4404. A single, well-formed colony from the selection plate was transferred to 50 ml of YEP selective medium that contained 50 mg/l kanamycin and maintained in a shaking incubator at 28 °C for 18 – 24 hrs or the length of time needed to reach an OD_600_ of 0.6 - 0.7. The *Agrobacterium* suspension was centrifuged at 8000 rpm for 10 min at 20°C. The pellet was resuspended in 50 ml of 2% MSO medium (4.3 g/l MS salts, 100 mg/l myo-inositol, 0.4 mg/l thiamine, and 20 g/l sucrose) by vortexing.

### Agrobacterium-mediated transformation

Cotyledon explants were incubated in the *Agrobacterium*/2% MSO suspension for 5 min, transferred to a sterile paper towel to allow excess suspension to briefly drain, placed back onto the feeder plates with the adaxial sides down, and co-cultivated in the dark at 19 °C for 48 hrs. Explants, adaxial side up, were transferred to our standard plant regeneration selective medium designated 2ZK that contained 4.3 g/l MS salts, 100 mg/l myo-inositol, 1 ml/l Nitsch vitamins (1000x), 20 g/l sucrose, 2 mg/l trans-zeatin, 75 mg/l kanamycin, 300 mg/l timentin, and 5.2 g/l Agargel. One week later, the explants were transferred onto 2ZK medium containing IAA at either 0 mg/l, 0.01 mg/l, 0.05 mg/l, 0.1 mg/l, or 0.5 mg/l IAA.

After two weeks, explants were transferred onto 1ZK medium that contained 4.3 g/l MS salts, 100 mg/l myo-inositol, 1 ml/l Nitsch vitamins (1000x), 20 g/l sucrose, 1 mg/l trans-zeatin, 75 mg/l kanamycin, 300 mg/l timentin, 5.2 g/l Agargel, and IAA at either 0 mg/l, 0.01 mg/l, 0.05 mg/l, or 0.1 mg/l IAA, or 0.5 mg/l in plates or Magenta GA7 boxes depending upon the size of the shoots regenerating from the cotyledon explants.

When shoots were approximately 3 mm tall, they were excised from the cotyledon explants and transferred to selective rooting medium designated RMK (4.3 g/l MS salts, 1 ml/l Nitsch vitamins (1000x), 30 g/l sucrose, pH 6.0, 8 g/l Difco Bacto agar (Becton, Dickinson and Company, Franklin Lakes, NJ), 75 mg/l kanamycin, 300 mg/l timentin, and IAA at either 0 mg/l or 1 mg/l in Magenta GA7 boxes.

Unless otherwise noted, the pH of all media was adjusted to 5.8 before autoclaving. For all media, the trans-zeatin, IAA, kanamycin, and timentin were dispensed from filter sterilized stock solutions into autoclaved medium that was allowed to cool to 55^o^C. Cotyledon explant cultures were transferred to freshly prepared medium every two weeks.

### GUS histochemical assay

Histochemical assay of b-glucuronidase (GUS) activity was performed on leaves from putative transgenic and control (non-transformed) plants. Leaves were vacuum infiltrated for 20 – 30 min in buffer (0.8 g/l 5-bromo-4-chloro-3-indolyl-b-D-glucuronide (X-Gluc), 0.1 M Na_2_HPO_4_, 0.1 M NaH_2_PO_4_ phosphate, 10 mM ethylenediamine tetraacetic acid (EDTA), 1.6 mM potassium-ferricyanide and 1.6 mM potassium–ferrocyanide, 5% v/v Triton X-100, and 20% v/v methanol) before incubation at 37 °C overnight. The chlorophyll was removed from the leaves by 3 - 4 washes with 70% ethanol at room temperature. The leaves were examined with a Leica S8APO stereomicroscope outfitted with a digital camera.

### Polymerase chain reaction analysis

To confirm the presence of the neomycin phosphotransferase II selectable marker gene (*nptII*), DNA was extracted from leaves of putative transgenic lines and controls (non-transformed) with the Qiagen DNeasy plant mini kit (Hilden, Germany) as per the manufacturer’s instructions. Primers used to detect *nptII* were forward 5’-GGC TGG AGA GGC TAT TC-3’ and reverse 5’-GGA GGC GAT AGA AGG CG-3’. The diagnostic amplicon size expected with these primers is approximately 700 bp. The PCR program started with a one-step cycle of 2 min at 95^o^C, followed by 29 cycles of 30 s at 94^o^C, 45 s at 57^o^C, 50 s at 72^o^C, and a 10 min final extension at 72^o^C. DNA was separated and visualized by electrophoresis through a 1% agaraose, ethidium bromide-stained gel.

### Experimental Design

A total of 5 different experiments were performed. Three biological replicates were used for each IAA concentration in each experiment. A total of 750 cotyledon explants were used per IAA concentration investigated. The standard error was calculated.

## Results

### Optimization of IAA concentration for recovery of stable transgenic lines

After the co-cultivation period that followed infection with *Agrobacterium*, cotyledon explants were transferred to our standard selective plant regeneration medium designated 2ZK that contains 2 mg/l trans-zeatin as the only plant growth regulator. One week later, the explants were transferred to 2ZK supplemented with different IAA concentrations (0 mg/l, 0.01 mg/l, 0.05 mg/l, 0.1 mg/l, 0.5 mg/l) to determine if the addition of IAA would decrease the time from infection with *Agrobacterium* to recovery of stable transgenic lines. We continued to use this same series of IAA concentrations in the subsequent selective plant regeneration medium designated 1ZK. Medium supplemented with IAA resulted in shoots that were more fully developed earlier in the culture process as compared to medium without IAA (Fig. 1A). In Figure 1A, cotyledon cultures shown in a – e represent controls that were not infected with *Agrobacterium*. We observed that as the IAA concentration increased, the level of plant regeneration from the controls decreased (Fig. 1A, a – d). Cotyledon explants infected with *Agrobacterium* and cultured on medium containing IAA exhibited the same pattern of shoot development as the cotyledons not infected, in that we observed more well-developed shoots at an early stage of culture post infection (Fig. 11 f – i). For our standard method without IAA (Fig. 1A, e), the level of plant regeneration is significantly less in comparison with medium that contained IAA.

**Figure 1.**
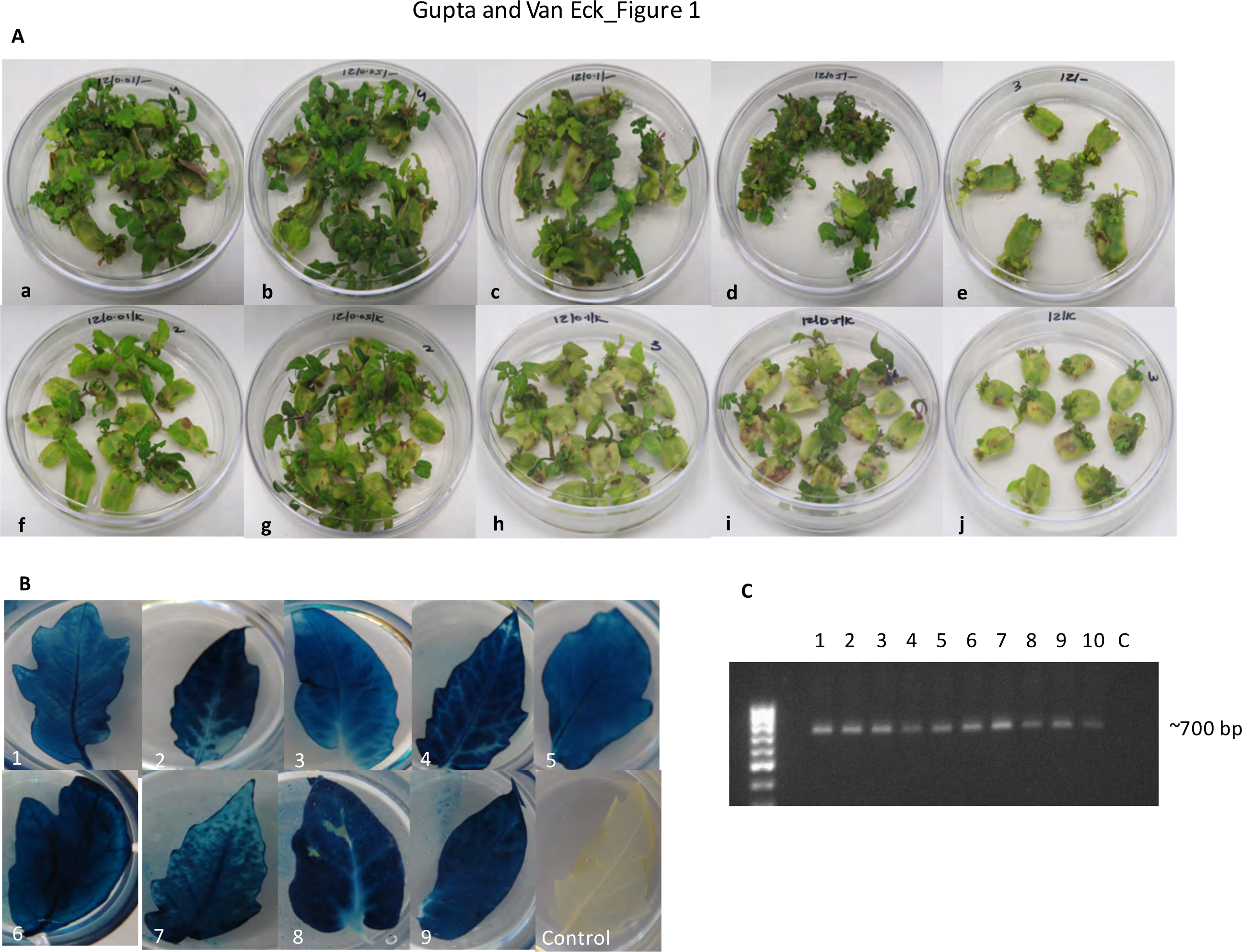
Results for the recovery of *Solanum lycopersicum* cv M82 stable transgenics from*Agrobacterium tumefaciens*-infected cotyledon explants cultured on plant regeneration medium supplemented with different concentrations of indole-3-acetic acid (IAA). (A) *Agrobacterium tumefaciens*-infected cotyledon explants (approximately 5 weeks post infection) cultured on selective plant regeneration medium containing the following amounts of IAA in mg/l (f) 0.01, (g) 0.05, (h) 0.1, (i) 0.5, and (j) 0. Images a – e represent the corresponding non-infected controls for each IAA concentration, respectively. (B) Histochemical analysis for GUS expression in leaves taken from independent transgenic lines designated 1 - 9 recovered from selective plant regeneration medium that contained 0.1 mg/l IAA. GUS expression was not observed in the non-transformed controls. (C) Agarose gel of PCR products showing the expected ∼700 bp product amplified from the *nptII* selectable marker gene in 10 independent transgenic lines (lanes 1 – 10). These lines were recovered from selective plant regeneration medium that contained 0.1 mg/l IAA. C = the control

In general, earlier emergence of well-developed shoots from *Agrobacterium*-infected cotyledon explants on medium containing IAA translated to the recovery of whole rooted plants in less time as compared with medium that did not contain IAA (Table 1). Medium containing either 0.05 mg/l or 0.1 mg/l IAA resulted in the shortest time, 11 wks, for recovery of stable transgenic lines. There appeared to be a threshold of IAA concentration and effect on recovery time because at 0.5 mg/l IAA the time was similar to our standard method. We observed a similar decrease in time when transformations of other tomato genotypes were performed by different lab members who tested plant regeneration medium that contained 0.1 mg/l IAA (data unpublished).

**Table 1.** Results for recovery of stable transgenic lines of *Solanum lycopersicum* cv M82 from *Agrobacterium tumefaciens*-infected cotyledon explants cultured on selective plant regeneration medium supplemented with different indole-3-acetic acid (IAA) concentrations. *Average transformation efficiency was calculated as percent of stable transgenic lines recovered from the total number of cotyledon explants infected with *Agrobacterium tumefaciens*. Transformation efficiency values shown are the average from 5 experiments ± the standard error (SE) calculated from 3 biological replicates.

| IAA<br>(mg/l) | Total number<br>explants | Total number<br>rooted plants | Average transformation<br>efficiency<br>(±) SE* | Total time for recovery of<br>transgenic lines<br>(wks) |
| --- | --- | --- | --- | --- |
| 0 | 750 | 660 | 88 ± 2.2 | 17 |
| 0.01 | 750 | 390 | 52 ± 1.0 | 15 |
| 0.05 | 750 | 375 | 50 ± 1.5 | 11 |
| 0.1 | 750 | 405 | 54 ± 1.2 | 11 |
| 0.5 | 750 | 360 | 48 ± 2.0 | 16 |

\*Average transformation efficiency was calculated as percent of stable transgenic lines recovered from the total number of cotyledon explants infected with *Agrobacterium tumefaciens*. Transformation efficiency values shown are the average from 5 experiments ± the standard error (SE) calculated from 3 biological replicates.

### Effect of IAA on transformation efficiency and rooting

The formula below was used to calculate transformation efficiency (TE):

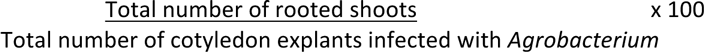

Overall, the TE was lower when medium containing IAA was used as compared to the TE of 88% when medium containing trans-zeatin as the only growth regulator was used (Table 1).

When putative transgenic lines were approximately 3 - 4 cm tall, they were removed from the cotyledon explants and transferred to either our standard selective rooting medium (RMK) without IAA or RMK supplemented with 1 mg/l IAA designated RMIK. We chose this concentration based on previous work with tomato transgenic lines recovered from a few genotypes that did not root as well as M82 on our standard rooting medium (data not published). We observed that shoots cultured on RMIK resulted in the emergence of roots after 6 - 7 days as compared to 11 - 14 days on RMK. The addition of IAA to the medium did not result in any phenotypic differences of the plants as compared to medium that did not contain IAA.

### Characterization of putative transgenic lines

The first level of analysis to confirm the recovered plants from *Agrobacterium*-infected cotyledons were transgenic was a histochemical assay for the GUS reporter protein. Whole leaves from plants rooted on RMK were used for the analysis. All leaves exhibited GUS activity, although we observed variation in the level of intensity with some leaves exhibiting a darker coloration than others (Fig 1B). GUS activity was not observed in leaves from non-transformed control plants.

To further confirm the recovered plants were indeed stable transgenic lines, we did PCR analysis for the presence of the *nptII* selectable marker gene in plants found to be positive for GUS activity. Total genomic DNA was isolated from the leaves of the GUS-positive lines and non-transformed control plants. PCR amplification of the *nptII* gene was detected in plants that were also GUS positive. No amplified product was detected in DNA from the control, (non-transgenic) plants (Fig. 1C).

### Modified protocol

Based on our findings, we now follow a modified protocol as outlined in Figure 2 for *Agrobacterium*-mediated tomato transformations. The modified protocol takes into account recovery time and TE. IAA concentrations of 0.05 and 0.1 mg/l IAA both resulted in a 6-week decrease for recovery of stable transformants, however, we chose to use 0.1 mg/l IAA in our modified protocol because of the 54% TE (Table 1). In addition to M82, we have applied this protocol to other tomato genotypes including the most closely related wild species, *Solanum pimpinellifolium*, and also observed a decrease in time for recovery of transgenic lines as compared to our previous tomato transformation methodology (data not published).

**Figure 2.**
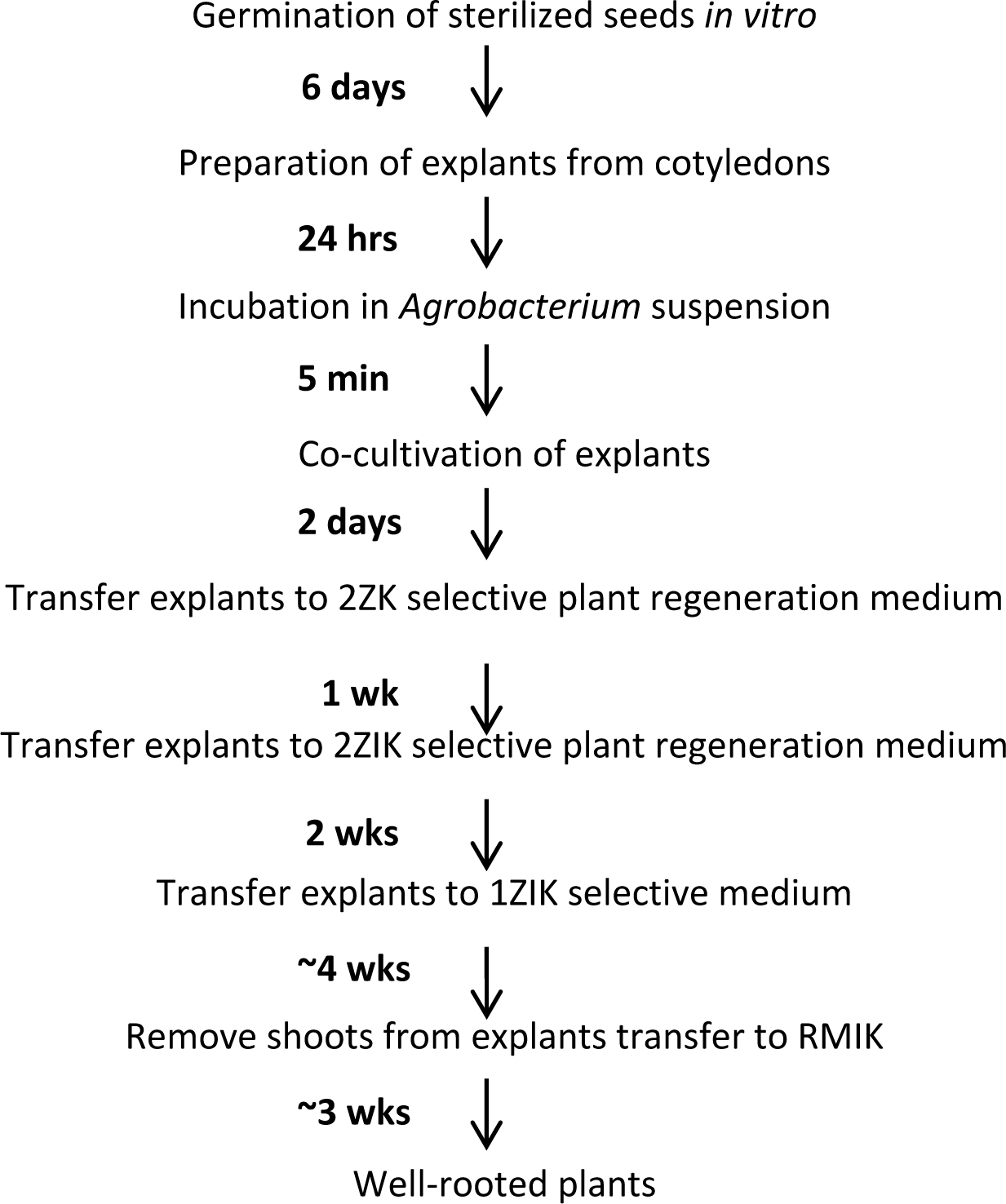
Schematic representation of the optimized *Agrobacterium tumefaciens*-mediated transformation methodology for *Solanum lycopersicum* cv M82. See the Materials and Methods for details on seed sterilization and all media compositions.

## Discussion

For development of stable transformation methodology, the foremost factors to be considered are transformation efficiency and the time from infection with *Agrobacterium tumefaciens* until the recovery of transgenic lines. Various parameters have been investigated to reach a high transformation efficiency for tomato including application of lipoic acid to reduce tissue necrosis caused by *Agrobacterium* infection of the MicroTom genotype (Dan et al. 2016). Methods that provide both high efficiency and the shortest time to recovery of transgenic lines lead to a high-throughput pipeline that allows earlier evaluation of gene function. In turn, a high-throughput pipeline decreases the amount of labor and resources needed, which can translate into significant financial savings.

The focus of our study was to investigate medium components that had the potential to decrease the time for recovery of stable tomato transgenic lines. Our standard method, which has a high transformation efficiency at approximately 90%, takes 17 weeks for recovery of transformants. The interest in optimization of our methods stemmed from an increased need for transgenic lines because tomato has become the model species of choice for many studies that include ripening, abiotic and biotic tolerance, and nutritional content (Gonzali et al. 2009; Martel et al. 2011; Nguyen et al. 2010; Sun et al. 2010). In addition, with the recent demonstration of successful genome editing by CRISPR/Cas9 in tomato, the interest in applying this technology for the study of gene function will increase (Brooks et al. 2014; Ito et al. 2015). Therefore, a transformation methodology that can deliver modified lines in a shorter time frame will help to advance these studies.

Our standard protocol is a modified version of methods reported by Fillatti et al. (1987) in which zeatin is the only growth regulator incorporated into the plant regeneration medium (Van Eck et al. 2006). We chose to start our investigation by examining additional growth regulators that, in combination with zeatin, would greatly reduce the time for recovery of stable transgenic lines but not have a significant negative effect on transformation efficiency. In a literature search, we found several reports that demonstrated a positive effect on tomato plant regeneration and transformation efficiency when indole-3-acetic acid (IAA) was incorporated into zeatin-containing plant regeneration medium (Gubis et al. 2004; Park et al. 2003; Yasmeen 2009). However, they did not report any effects observed on the time required to recover transgenic plants.

We found that addition of either 0.05 or 0.1 mg/l IAA to our standard plant regeneration medium that contains trans-zeatin as the only growth regulator decreased the time for recovery of stable transgenic lines from 17 to 11 weeks. Previous reports have demonstrated that shoot apical meristem development involves interactions among cytokinin signaling pathway components, auxin, and several families of transcription factors ( 2012). It is possible that the addition of IAA to our standard plant regeneration medium facilitates interactions among the cytokinin signaling and auxin regulated genes, which results in faster shoot development from the cotyledon explants.

Although there was a reduction in transformation efficiency with the addition of IAA from approximately 90% to about 50%, this level is acceptable considering transgenic lines can be evaluated significantly earlier than when our standard method was used. This decrease in time allows researchers to test their material earlier and make changes to their approaches sooner if results are unsatisfactory for their genes of interest.

In addition to supplementation of the standard plant regeneration medium with IAA, we also investigated effects of adding IAA to the rooting medium, which was not a component in our standard rooting medium. Inclusion of IAA in *in vitro* rooting medium has been reported for tomato, however, it is not routinely added because tomato readily develops roots in culture medium without growth regulators (Frary and Earle 1996). Our interest was to determine if supplementation decreased the time to rooting, which we did observe. Auxin is produced in both shoots and roots and the auxin produced in the roots helps in root development (Overvoorde et al. 2010; Petersson et al. 2009; Stepanova et al. 2008). It is possible that IAA, when exogenously added, increases the levels of auxin in the plants, hence resulting in the cells differentiating earlier to form roots. However, research needs to be conducted to confirm this hypothesis.

## Conclusions

Interest in tomato as a model has increased over the years and we have seen a rise in the number of research groups that require stable transgenic lines for various studies. Modification of our standard plant regeneration medium through the addition of either 0.05 or 0.1 mg/l AA shortened the recovery of transgenic lines by 6 weeks for the M82 tomato cultivar. Application of this modification for transformation of other tomato genotypes in our lab also resulted in a decreased time for recovery of stable transgenic lines.

A shorter recovery time for stable transgenic lines is highly desirable for functional studies to allow earlier determination of the genes and networks involved in phenotypes of interest. A decrease in recovery time would also provide a higher throughput process, which has the potential for cost savings related to labor and resources. Optimization studies of standard transformation methodologies for different plant species should always be considered in order to alleviate bottlenecks for generation of stable transgenic lines (Altpeter et al. 2016). Availability of efficient transformation methods is especially critical with the rapid development of genome editing technologies, which will result in an increased demand for generation of transgenic lines for basic research studies that can lead to crop improvement.

## Acknowledgements

We thank Cynthia Du for assistance with part of the experimental process. We thank Cynthia Du, Patricia Keen, and Michelle Tjahjadi for their critical review of the manuscript. Support for this work was through a grant from the National Science Foundation Plant Genome Research Program (IOS-1237880).

## Author contribution statement

SG and JVE designed the experiments, SG performed the experiments, SG and JVE wrote the manuscript. Both authors read and approved the manuscript.

## Compliance with ethical standards

### Conflict of interest

The authors declare that they have no competing interests.

Key Message: We decreased the time for recovery of tomato transgenic lines by 6 weeks through the addition of indole-3-acetic acid to our standard plant regeneration medium.

## References

Altpeter F, Springer NM, Bartley LE, Blechl AE, Brutnell TP, Citovsky V, Conrad LJ, Gelvin SB, Jackson DP, Kausch AP, Lemaux PG, Medford JI, Orozco-Cárdenas ML, Tricoli DM, Van Eck J, Voytas DF, Walbot V, Wang K, Zhang ZJ, Stewart CN, Jr. (2016) Advancing crop transformation in the era of genome editing. Plant Cell 28:1510–1520

Bombarely A, Menda N, Tecle IY, Buels RM, Strickler S, Fischer-York T, Pujar A, Leto J, Gosselin J, Mueller LA (2011) The Sol Genomics Network (solgenomics.net): growing tomatoes using Perl. Nucleic Acids Res 39:1149–1155

Brooks C, Nekrasov V, Lippman ZB, Van Eck J(2014) Efficient gene editing in tomato in the first generation using the Clustered Regularly Interspaced Short Palindromic Repeats/CRISPR-Associated9 system. Plant Physiol 166:1292–1297

Chen P-Y, Wang C-K, Soong S-C, To K-Y (2003) Complete sequence of the binary vector pBI121 and its application in cloning T-DNA insertion from transgenic plants. Mol Breeding 11: 287–293.

Chyi Y-S, Phillips GC (1987) High efficiency Agrobacterium-mediated transformation of Lycopersicon based on conditions favorable for regeneration. Plant Cell Rep 6:105–108

Consortium TTG (2012) The tomato genome sequence provides insights into fleshy fruit evolution. Nature 485:635–641

Dan Y, Zhang S, Matherly A(2016) Regulatoin of hydrogen peroxide accumulation and death of Agrobacterium-transformed cells in tomato transformation. Plant Cell Tiss Organ Cult doi:10.1007/s11240-016-1045-y

Emmanuel E, Levy AA (2002) Tomato mutants as tools for functional genomics. Curr Opin Plant Biol 5:112–117

Fillatti JJ, Kiser J, Rose R, Comai L(1987) Efficient transfer of a glyphosate tolerance gene into tomato using a binary Agrobacterium tumefaciens vector. Bio-Technol 5:726–730

Frary A, Earle ED (1996) An examination of factors affecting the efficiency of Agrobacterium-mediated transformation of tomato. Plant Cell Rep 16:235–240

Friml J, Vieten A, Sauer M, Weijers D, Schwarz H, Hamann T, Offringa R, Jurgens G(2003) Efflux-dependent auxin gradients establish the apical-basal axis of Arabidopsis. Nature 426:147–153

Gelvin SB(2003) Agrobacterium-mediated plant transformation: the biology behind the ” gene-jockeying” tool. Microbiol Mol Biol Rev 67:16–37

Gonzali S, Mazzucato A, Perata P (2009) Purple as a tomato: towards high anthocyanin tomatoes. Trends Plant Sci 14:237–241

Gubis J, Lajchova Z, Farago J, Jurekova Z(2004) Effect of growth regulators on shoot induction and plant regeneration in tomato (Lycopersicon esculentum Mill.). Biologia 59:405–408

Gupta S, Rashotte AM (2012) Down-stream components of cytokinin signaling and the role of cytokinin throughout the plant. Plant Cell Rep 31:801–812

Ito Y, Nishizawa-Yokoi A, Endo M, Mikami M, Toki S (2015) CRISPR/Cas9-mediated mutagenesis of the RIN locus that regulates tomato fruit ripening. Biochem Biophys Res Commun 467:76–82

Jones JBJ(1998) Tomato Plant Culture: In the Field, Greenhouse, and Home Garden. CRC Press LLC, Boca Raton, FL

Martel C, Vrebalov J, Tafelmeyer P, Giovannoni JJ (2011) The tomato MADS-box transcription factor RIPENING INHIBITOR interacts with promoters involved in numerous ripening processes in a COLORLESS NONRIPENING-dependent manner. Plant Physiol 157:1568–1579

Mccormick S, Niedermeyer J, Fry J, Barnason A, Horsch R, Fraley R (1986) Leaf disk transformation of cultivated tomato (L. esculentum) using Agrobacterium tumefaciens. x Plant Cell Rep 5:81–84

Murashige T, Skoog F (1962) A revised medium for rapid growth and bio assays with tobacco tissue cultures. Physiol Plantarum 15:473–497

Nguyen HP, Chakravarthy S, Velasquez AC, McLane HL, Zeng L, Nakayashiki H, Park DH, Collmer A, Martin GB (2010) Methods to study PAMP-triggered immunity using tomato and Nicotiana benthamiana. Mol Plant Microbe Interact 23:991–999

Overvoorde P, Fukaki H, Beeckman T (2010) Auxin control of root development. Cold Spring Harb Perspect Biol 21: 1–16.

Park SH, Morris JL, Park JE, Hirschi KD, Smith RH(2003) Efficient and genotype-independent Agrobacterium-mediated tomato transformation. J Plant Physiol 160:1253–1257

Petersson SV, Johansson AI, Kowalczyk M, Makoveychuk A, Wang JY, Moritz T, Grebe M, Benfey PN, Sandberg G, Ljung K (2009) An auxin gradient and maximum in the Arabidopsis root apex shown by high-resolution cell-specific analysis of IAA distribution and synthesis. Plant Cell 21:1659–1668

Pitzschke A, Hirt H (2010) New insights into an old story:Agrobacterium-induced tumour formation in plants by plant transformation. EMBO J 29:1021–1032

Sabatini S, Beis D, Wolkenfelt H, Murfett J, Guilfoyle T, Malamy J, Benfey P, Leyser O, Bechtold N, Weisbeek P, Scheres B (1999) An auxin-dependent distal organizer of pattern and polarity in the Arabidopsis root. Cell 99:463–472

Shimizu-Sato S, Tanaka M, Mori H (2009) Auxin-cytokinin interactions in the control of shoot branching. Plant Mol Biol 69:429–435

Somerville C, Koornneef M (2002) A fortunate choice: the history of Arabidopsis as a model plant. Nat Rev Genet 3:883–889

Stepanova AN, Robertson-Hoyt J, Yun J, Benavente LM, Xie DY, DoleZal K, Schlereth A, Jurgens G, Alonso JM (2008) TAA1-mediated auxin biosynthesis is essential for hormone crosstalk and plant development. Cell 133:177–191

Sun HJ, Uchii S, Watanabe S, Ezura H(2006) A highly efficient transformation protocol for Micro-Tom, a model cultivar for tomato functional genomics. Plant Cell Physiol 47:426–431

Sun W, Xu X, Zhu H, Liu A, Liu L, Li J, Hua X(2010) Comparative transcriptomic profiling of a salt-tolerant wild tomato species and a salt-sensitive tomato cultivar. Plant Cell Physiol 51:997–1006

Van Eck J, Kirk DD, Walmsley AM (2006) Tomato (Lycopersicum esculentum). In: Wang K (ed) Methods in Molecular Biology, Agrobacterium Protocols. vol 343. Humana Press Inc., Totowa, NJ, pp 459–473

Xu C, Liberatore KL, MacAlister CA, Huang Z, Chu YH, Jiang K, Brooks C, Ogawa-Ohnishi M, Xiong G, Pauly M, Van Eck J, Matsubayashi Y, van der Knaap E, Lippman ZB (2015) A cascade of arabinosyltransferases controls shoot meristem size in tomato. Nat Genet 47:784–792

Yasmeen A(2009) An improved protocol for the regeneration and transformation of tomato (cv Rio Grande). Acta Physiol Plant 31:1271–1277

